# Comparison of antioxidant activity between cyanidin-3-O-glucoside (C3G) liposome and cyanidin-3-O-glucoside (C3G) in 2D and 3D cell cultures

**DOI:** 10.1101/314369

**Authors:** Tisong Liang, Rongfa Guan, Guozhou Cao, Haitao Shen, Zhenfeng Liu, Qile Xia, Zhe Wang

**Affiliations:** Zhejiang Provincial Key Laboratory of Biometrology and Inspection and Quarantine, China Jiliang University, Hangzhou 310018, China; Ningbo Academy of Inspection and Quarantine, ningbo yingyi road No.66 A building room 518, nignbo 315012,China; Zhejiang Provincial Center for Disease Control and Prevention, 3399 Binsheng Road, Hangzhou 310051, China; Chiatai Qingchunbao Pharmaceutical Co., LTD, NO. 551 Xixi Road, Hangzhou 310023, China; Food Science Institute, Zhejiang Academy of Agricultural Sciences, 298 Desheng Road, Hangzhou 310021, China

**Keywords:** 3D cell culture, C3G, C3G liposomes, Antioxidant activity, Caco-2 cell

## Abstract

The 2D cell culture is the predominant in vitro model for numerous studies. However, 2D cell cultures may not accurately reflect the functions of three-dimensional (3D) tissues, which have extensive cell–cell and cell–matrix interactions; thus, using 2D cell cultures may lead to inaccurate experimental results. Therefore, to obtain adequate and detailed information about the antioxidant activity of cyanidin-3-O-glucoside (C3G) and C3G liposomes in the 2D and 3D cell culture models, we used in this study H_2_O_2_ to construct the cell damage model and assess the antioxidant activity of C3G and C3G liposomes on Caco-2 cells cultured in the 3D model. We also measured the cell viability, cell morphology, and activity of glutathione (GSH), superoxide dismutase (SOD), total antioxidant capacity (T-AOC), and malondialdehyde (MDA) content of Caco-2 cells treated with H_2_O_2_, C3G, and C3G liposomes. Results showed that cells cultured in the 3D culture model formed a 3D structure and tight spheroids and showed increased cell activity and IC_50_. The C3G and C3G liposomes can enhance the activity of GSH, SOD, and T-AOC but decrease the MDA content. At the same time, the effect was more obvious in the 3D cell culture model than in the cells cultured in the 2D model. This study revealed that the results obtained from the 2D cell model may be inaccurate compared with the results obtained from the 3D cell model. A realistic mechanism study of antioxidant activity of C3G and C3G liposomes in the 3D cell model, which acts as an intermediate stage bridging the in vitro 2D and in vivo models, was observed.

## 1. Introduction

Anthocyanins are water-soluble vacuolar pigments found in most species in the plant kingdom and are responsible for the red, purple, and blue colors of many fruits, vegetables, and flowers^1^, ^2^. Anthocyanins exhibit potential free radical-scavenging activities that prevent low-density lipoprotein oxidation and positively affect chronic gut inflammatory diseases, obesity, and inflammation^3-5^. Moreover, cyanidin-3-O-glucoside (C3G), which is an anthocyanin with a high percentage and belongs to the flavonoid family and commonly present in human diet, exhibits antiinflammatory and antioxidant effects^6-8^. Some studies have showed that C3G protects from the adverse effects of radiation, controls the key aspects of tumorigenesis, inhibits proliferation, induces apoptosis of cancer cells, reduces oxidative stress, fights H_2_O_2_-induced oxidative stress in human embryonic kidney cells, induces cell apoptosis and inhibit cell migration^9-15^.

Liposomes are vesicles, wherein the small aqueous volumes are surrounded by bilayer membranes that are normally composed of phospholipids^16^, ^17^. Liposomes can enhance the stability and bioavailability of encapsulated materials by protecting them from external environment factors, making them ideal candidates for drug delivery and important in food systems^18-21^.

The two-dimensional (2D) monolayer cell models have low anatomical and physiological relevance and remain the predominant in vitro model for numerous studies^22^, ^23^. However, some disadvantages exist in the use of the 2D monolayer cell model in vitro in some studies. Growing concerns have been demonstrating that monolayer and monotypic (2D) cellular screening assays may not effectively reproduce the response of a three-dimensional (3D) solid tumor to pharmacological compounds^24^, ^25^. In cancer research, the limitations of the 2D models are considered one major reason for the approximately 95% of potential anticancer drugs failures in clinical trials ^26^.

The 3D spheroid cell culture systems are providing new insights into tumor biology and also in differentiation, tissue organization, and homeostasis^27^, ^28^. Cells in in vitro 3D culture systems are in league with their counterparts in vivo compared with cells grown as 2D monolayers, wherein 2D monolayer cells also exhibit altered cell cycle durations, morphologies, susceptibility to drugs, metabolism, and gene and protein expression levels^29^, ^30^. Cells cultured as 3D models exhibit features that are close to the complex in vivo conditions^31^ and have been proven realistic for translating the study findings for in vivo applications; moreover, culturing cell lines as 3D models induces them to behave a step closer to the natural conditions^32^, ^33^. The 3D models also can help investigate the interplay between different physiological conditions (oxygen or nutrient deprivation), irradiation, or other physical and chemical stimuli^34^.We used liposome technology because liposomes can protect embedding materials and thus improve the effectiveness and stability of C3G, which could be used as a carrier.

Therefore, the objectives of this study were to compare the antioxidant activity of C3G and C3G liposomes by assessing the cellular viability in Caco-2 cells cultured in 2D and 3D conditions. We compared the antioxidant activity via the assay of glutathione (GSH), superoxide dismutase (SOD), malondialdehyde (MDA), and total antioxidant capacity (T-AOC).

## 2. Materials and methods

### 2.1 Materials

The Caco-2 cells (CBCAS, Shanghai, China), Cyanidin-3-glucoside (C3G) was purchased from Chengdu Biopurify Phytochemicals Ltd (Chengdu,China). Phosphatidylcholine (PC) and cholesterol (CH) were purchased from Beijing Shuangxuan Microorganism Co. Ltd (Beijing, China). Chloroform and diethyl ether were obtained from Hangzhou Jiachen Chemical Company (Hangzhou, China). All other chemicals were of reagent grade. The water used for all experiments was deionized water.

### 2.2 Methods

#### 2.2.1 Preparation of C3G liposomes

C3G liposomes were prepared by reverse-phase evaporation method^35^, ^36^. Phosphatidylcholine and cholesterol were dissolved in chloroform-diethyl ether(W_PC_: W_CH_=2.87:1). C3G was dissolved in phosphate-buffered saline (PBS) (0.20 M, pH 7.4). Then, the organic phase was homogenized with the aqueous phase by probe sonication for 10 min. The mixture was transferred to a round-bottomed flask. The organic solvent was evaporated under reduced pressure with a rotary evaporator to form a gel. Subsequently, 30 mL of phosphate-buffered solution was added to the gel, which was then probe-sonicated for an additional 25 min. The as-prepared liposomes were stored at 4 °C for further study.

#### 2.2.2 Morphology of the liposomes

Transmission electron microscopy (JEM-2100, Japanese Electronics Co., Ltd., Tokyo, Japan) was employed to determine the microstructure of C3G liposomes via negative staining method. A drop of this solution was placed on a Formvar-carbon-coated copper grid for 5 min and was then imaged^37^.

#### 2.2.3 Cell culture

##### 2.2.3.1 Cell culture

The Caco-2 cells (CBCAS, Shanghai, China) were maintained in minimum essential medium (MEM) (Hyclone Laboratories, Inc., USA) supplemented with 10% fetal bovine serum (FBS) Hyclone Laboratories, Inc., USA, penicillin (100 kU/L), and streptomycin (100 g/L) at 37 °C under a 5% CO_2_ atmosphere in a humidified incubator ^38-40^. After reaching 70%–80% confluence, the cells were subcultured and maintained with medium changes every 2–3 days.

##### 2.2.3.2 2D cell culture

Caco-2 cells were isolated using 0.25% Trypsin-EDTA (Sigma, USA) and centrifuged at 1000 rpm for 5 min. Isolated cells were seeded at 24- and 96-well plates according to the experimental design. The cells were then cultured using MEM supplemented with 10% FBS (Gibco BRL Co., Ltd., USA) and 1% antibiotics (100 IU/mL penicillin, and 100 μg/mL streptomycin) and maintained at 37 °C in an atmosphere of 95% air with 5% CO_2_. All the experiments were performed on logarithmically growing cells. Prepared cells were exposed to various C3G liposome concentrations (0, 0.05, 0.10, 0.15, 0.20, or 0.25 mg/mL).

##### 2.2.3.3 3D cell culture

The 3D cell culture was carried out using the 3D cell culture hydrogel kit (Hangzhou Kevin Biotechnology Co., Ltd., Hangzhou, China) in accordance with the manufacturer’s directions. The Caco-2 cells were isolated as previously described, and the cell suspension was mixed with the gel (vol:vol=1:1) and shaken slightly and rapidly on a vortex for 1 s and repeated 1–2 times. Subsequently, the gel–cell mixture was added to the well and incubated in the plate in an incubator at 37 °C for 5–10 min. Lastly, the culture medium was added to the plate, and the plate was moved to the humidified incubator and maintained at 37 °C in an atmosphere with 5% CO_2_. The medium was replaced every 1–2 days. All the experiments were performed on logarithmically growing cells.

#### 2.2.4 Cell morphology

For documenting the morphology of Caco-2 cells in 3D culture condition, the cells were washed with PBS twice. Phase contrast images of the cells cultured in 3D conditions were observed using an inverted microscope (Nikon Eclipse Ti, Nikon, Tokyo, Japan) at different magnifications^41^.

Scanning electron microscopy was used to analyze the cell morphology in different cell culture models. The Caco-2 cells (1×10^5^) were seeded on glass slides placed in a 6-well plate for 2 days. Then, the samples were removed from culture wells, rinsed with PBS, and fixed with 2.5% glutaraldehyde for 8 h at 4 °C. Subsequently, the samples were rinsed again and postfixed in osmium tetroxide for 1 h before being dehydrated in a series of graded ethanol solutions (30%, 50%, 70%, 80%, 90%, and 95% vol/vol). Final drying was performed by critical-point method. Finally, gold-coated specimens were examined by SEM (Hitachi, SU-8010, Tokyo, Japan)^42^.

#### 2.2.5 Cell viability

Cell viability was determined by 2,5-diphenyl tetrazolium bromide (MTT) assay at 550 nm with a microplate reader (Tecan Co., Weymouth, UK)^43^. The cells were seeded in 96-well plates in a 100 μL volume (1 × 10^4^ per well) and allowed to grow for 24 h before treatment with different concentrations of C3G liposomes for 24 h. At the end of the experiments, 10 μL of 5 μg/mL MTT was added to each well. The cells were then incubated at 37 °C for 4 h. Formazan was solubilized in 150 μL of dimethyl sulfoxide and measured at 550 nm. The results were given as relative values to the negative control in percentage, whereas the positive control was set to be 100% viable.

The percentage of cell proliferation was calculated as^44^:

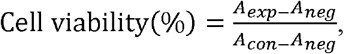

where *A_exp_* is the amount of experimental group absorbance, *A_neg_* is the amount of blank group absorbance, and *A_con_* is the amount of the control group absorbance.

#### 2.2.6 Assay of GSH and SOD activity

The assay of GSH and SOD activities was detected by glutathione peroxidase assay kit and total superoxide dismutase assay kit according to the manufacturer’s protocol (Nanjing Jiancheng Bioengineering Institute, China). The Caco-2 cells were cultured in a 25 cm^2^ culture flask and exposed to C3G and C3G liposomes for 12 h. The cells were then collected through centrifugation at 400 ×*g* for 5 min at 4 °C and the supernatant was removed. The cell suspension was washed and centrifuged twice using ice-cold PBS to remove all traces of the medium. The cell pellet was sonicated at 300 W for 10 s (three cycles) to obtain lysates, which were then centrifuged at 12,000 rpm for 10 min. Cell supernatants were finally assayed to measure GSH and SOD activities^45,46^.

Exactly 600 μL of the cell suspension, 150 μL of the substrate solution, and 600 μL of the reaction buffer solution were transferred to a fresh tube. The standard group included 25 μmol/L GSH dissolved in a GSH buffer solution. In the blank group, GSH was replaced by PBS. A microplate reader estimated the absorbance at 405 nm, and protein content was measured using the Bradford method using bovine serum albumin as the standard.

Exactly 20 μL of the prepared sample, 200 μL of the SOD working solution, 20 μL of the enzyme working solution, and 20 μL of dilution buffer were added to each well of a 96-well plate, and the mixture was mixed thoroughly. The plate was incubated for 20 min at 37 °C, and a plate reader was used to detect the absorbance at 450 nm.

#### 2.2.7 T-AOC assay and MDA assay

The Caco-2 cells were cultured in a 25 cm^2^ culture flask and exposed to C3G and C3G liposomes for 12 h and collected for the T-AOC and MDA level tests. The T-AOC and MDA levels in Caco-2 cells were measured with T-AOC and MDA analysis kits (Nanjing Jiancheng Bioengineering Institute, Nanjing, China) following the manufacturer’s protocols^47^, ^48^.

The Caco-2 cells were cultured in a 25 cm^2^ culture flask and exposed to C3G and C3G liposomes for 12 h. The cells were then collected through centrifugation at 400×*g* for 5 min at 4 °C and the supernatant was removed. The cell suspension was washed and centrifuged twice using ice-cold PBS to remove all traces of the medium. The cell pellet was sonicated at 300 W for 10 s (three cycles) to obtain lysates, which were then centrifuged at 12,000 rpm for 10 min. Cell supernatants were finally assayed to measure the assay of T-AOC and MDA.

For the detected of T-AOC of cells, exactly 100 μL of the cell suspension added the reagent and the samples were treated according the manufacturer’s protocols. A microplate reader estimated the absorbance at 520 nm, and protein content was measured using the Bradford method using bovine serum albumin as the standard.

For the detected of MDA, exactly 200μL of the cell suspension added the reagent and the samples were treated according the manufacturer’s protocols.The plate was incubated for 20 min at 37 °C, and a plate reader was used to detect the absorbance at 532nm.

#### 2.2.8 Statistical analysis

All experiments were performed thrice in duplicate per sample. Data are presented as the means ± standard deviations from at least three independent measurements. Statistical analysis was performed using SPSS version 21.0 for Windows.

## 3 Results

### 3.1 Characterization studies

Transmission electron microscopy (TEM) was used to investigate the morphology of C3G liposomes. Figure 1 shows a recorded representative TEM image of C3G liposomes. The nanoparticles exhibited spherical shapes, and the size of C3G liposomes was approximately 200 nm and formed a vesicular structure. From our previous study^36^,the average diameter of C3G liposomes is 165.78±4.3nm, the encapsulation efficiency of C3G liposomes is 70.43%±1.95%.

**Fig. 1.**
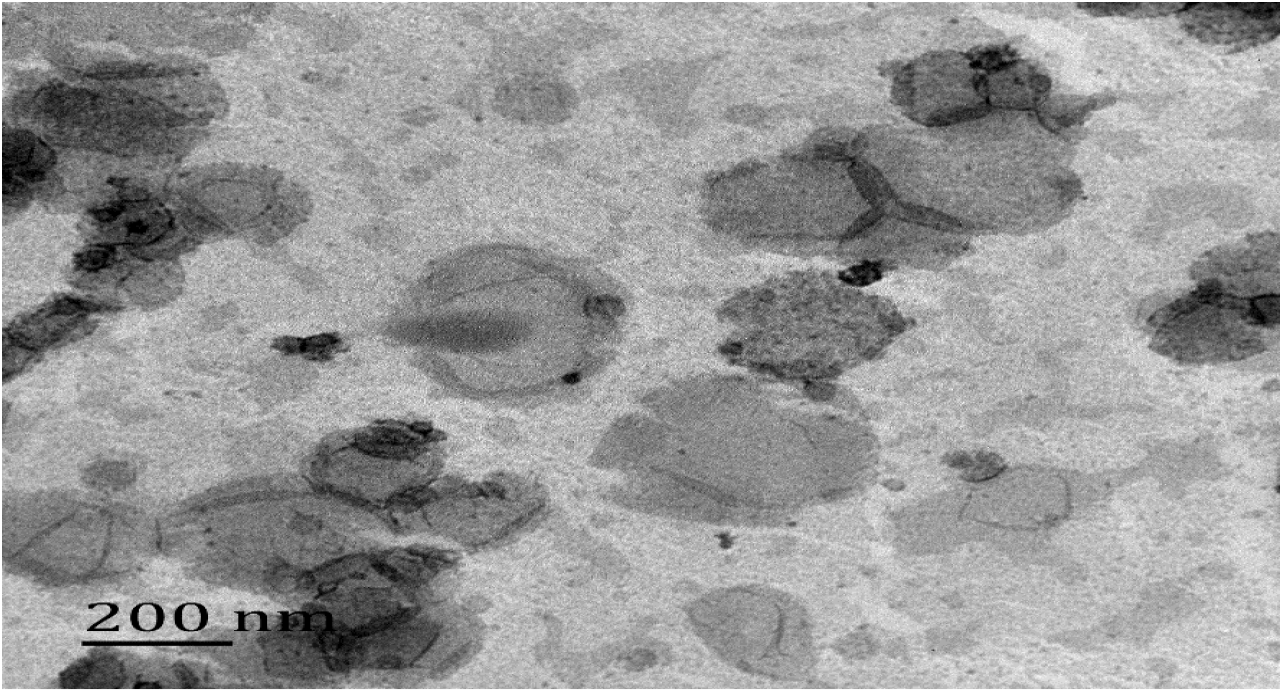
Transmission Election Microscope (TEM) image of C3G nanoliposomes.

### 3.2 Cell morphology

In our work, Caco-2 cells were cultured in the 3D models for 7 days and washed with PBS thrice. Subsequently, cell morphology was observed at different magnifications using an inverted microscope and scanning electron microscope.

As shown in Fig.2 and Fig.3, the morphology of the cell culture in the 3D model was different from the cell culture in the 2D model. From Fig.2, at different magnifications, the cells showed a mass-like spheroid growth, and the cells gathered and grew together. From the scanning electron microscope image, the Caco-2 cells grew together in patches in the 2D culture. However, when grown in the 3D cell culture model, they formed uniform spheroids with relatively smooth surfaces. The SEM image showed that the Caco-2 cell culture in the 3D culture model formed a 3D structure and tight spheroids.

**Fig. 2.**
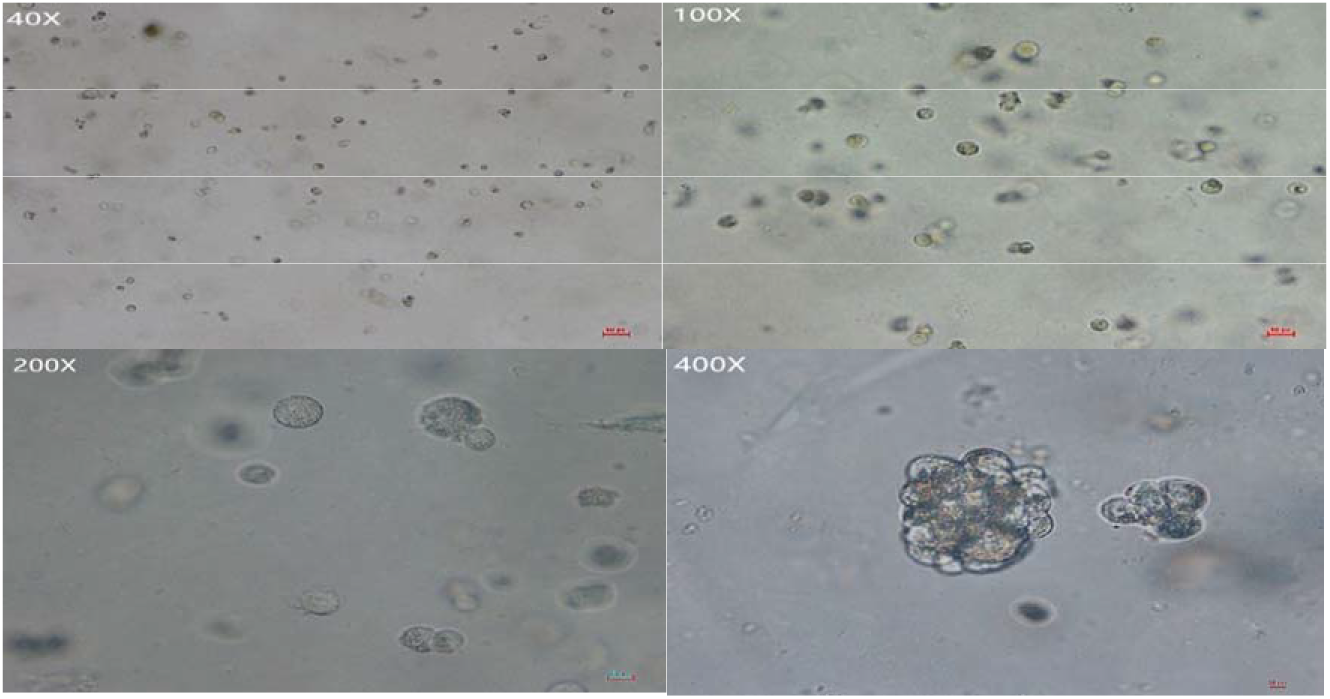
Cell morphology of 3D culture cells in different magnification times.

**Fig. 3.**
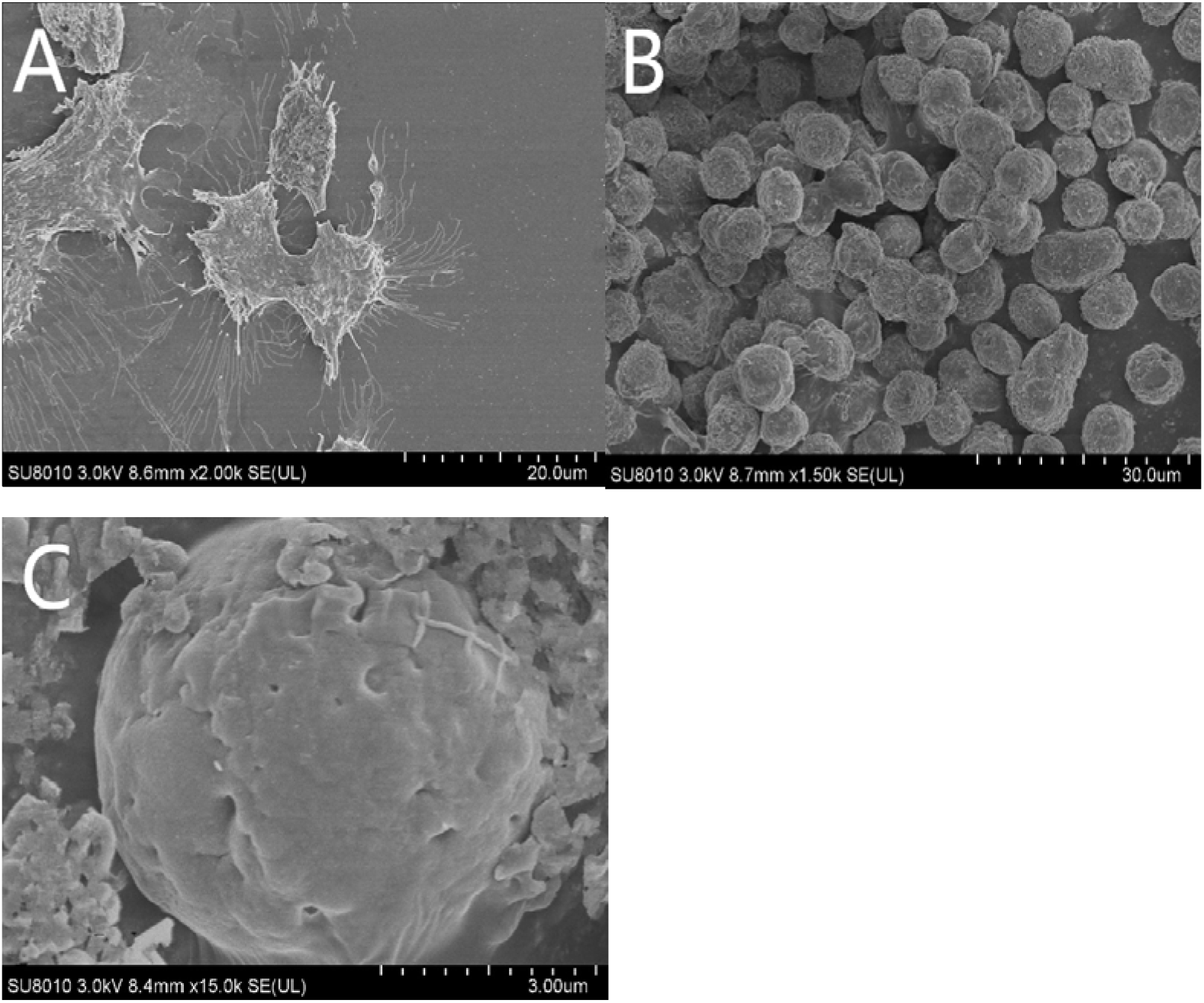
The cell morphology of 2D and 3D cell culture models(A:The cells culture in 2D cell culture model;B:The cells culture in 2D cell culture model that treated with pancreatin;C: The cells culture in 3D cell culture model.)

### 3.3 Cell viability

In this study, Caco-2 cells cultured in 2D and 3D models were treated with C3G and C3G liposomes. Then, the cell viability was detected by MTT assay. The result is shown in Fig.4.

**Fig. 4.**
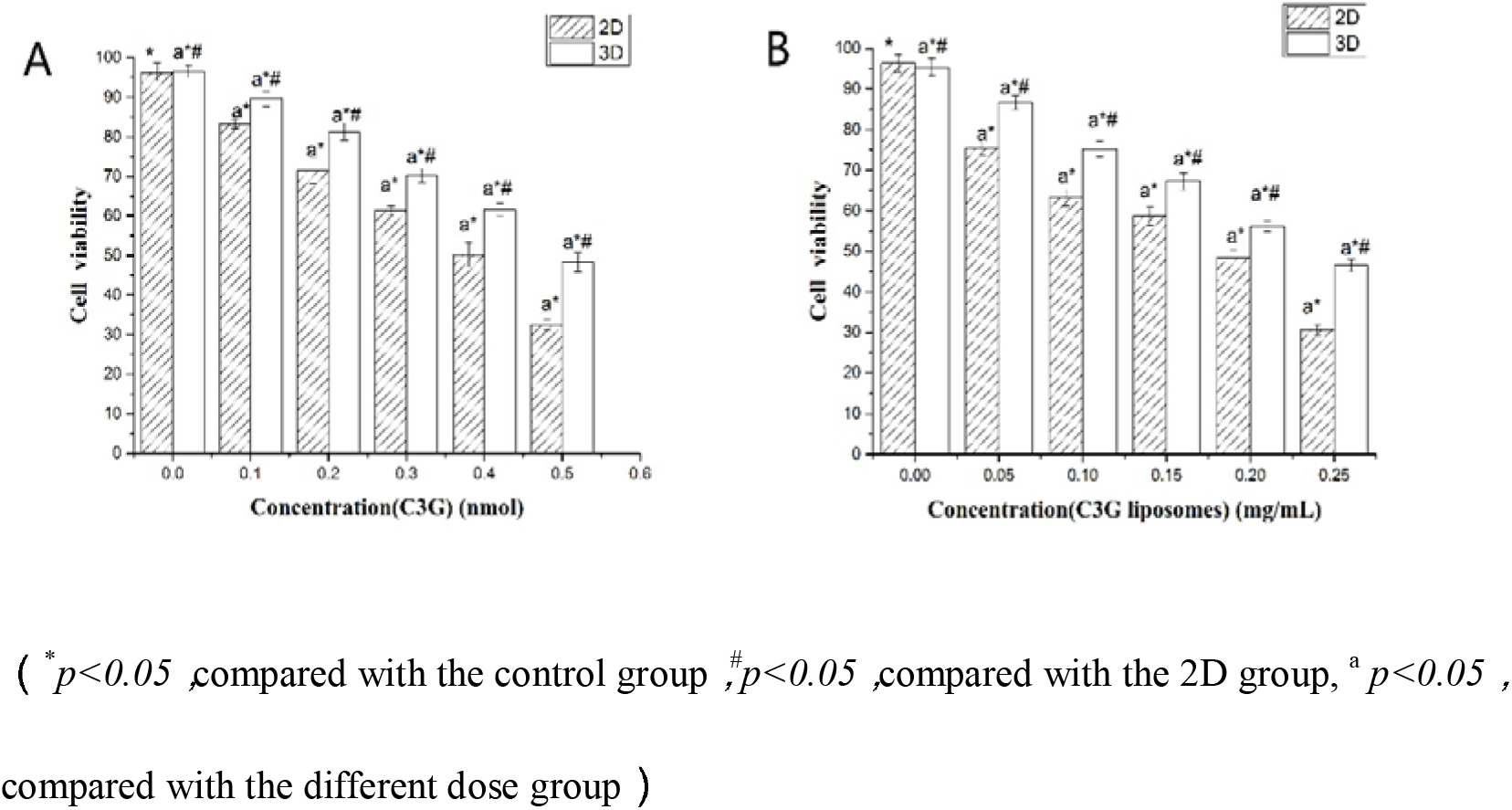
The cell viability of 2D and 3D culture cells expose to C3G and C3G liposomes (A:The cell treated with C3G;B:The cell treated with C3G liposomes.)

As shown in Fig. 4, the MTT assay results demonstrated a concentration-dependent activity after exposure to C3G and C3G liposomes in different cell culture models. From Fig. 4A, the cells in different culture models exhibit different cell activities after being exposed to C3G. In the 3D culture model, the cell activity is higher than that cultured in the 2D culture model. IC_50_ of cells exposed to C3G in the 2D and 3D cell culture models are 0.19 and 0.25mg/mL, and 0.175 and 0.233 mg/mL when exposed to C3G liposomes in different culture models, respectively. Figure 4B shows that the activity of cells exposed to C3G liposomes and cells cultured in the 3D culture model is higher than the cells cultured in the 2D culture model. Cells cultured in the 3D culture model show higher cell activity and IC_50_ compared with the 2D cell culture model in this study. This phenomenon is due to multicellular resistance and the adhesion between cells, thereby reducing the sensibility of Caco-2 cells to the C3G and C3G liposomes^49^, ^50^. At the same time, collagen gels purported to represent well in vivo conditions possibly causes the strong increase of cell viability.

### 3.4 Assay of GSH activity in different cell culture models

To compare the difference of GSH activity in different cell culture models, the Caco-2 cells cultured in the 2D and 3D culture models were treated with H_2_O_2_(0.2mmol), C3G, and C3G liposomes for 12 h, and the GSH activity was tested. Figure 5 shows the GSH activity in the 2D and 3D cell models. From Fig. 5, the GSH activity in different cell culture models is significantly different (*p*<0.05). The C3G and C3G liposomes both can enhance the GSH activity treated with H_2_O_2_, and the enhancement in the 3D model is higher than that in the 2D model. At the same time, the enhancement of C3G liposome is higher than that of C3G. This phenomenon is because the liposomes protect the C3G from the effect of external environment factors, leading to the high effectiveness of cells and multicellular resistance, and the adhesion between cells reduces the sensibility of Caco-2 cells to the C3G and C3G liposomes^49,50^

**Fig. 5.**
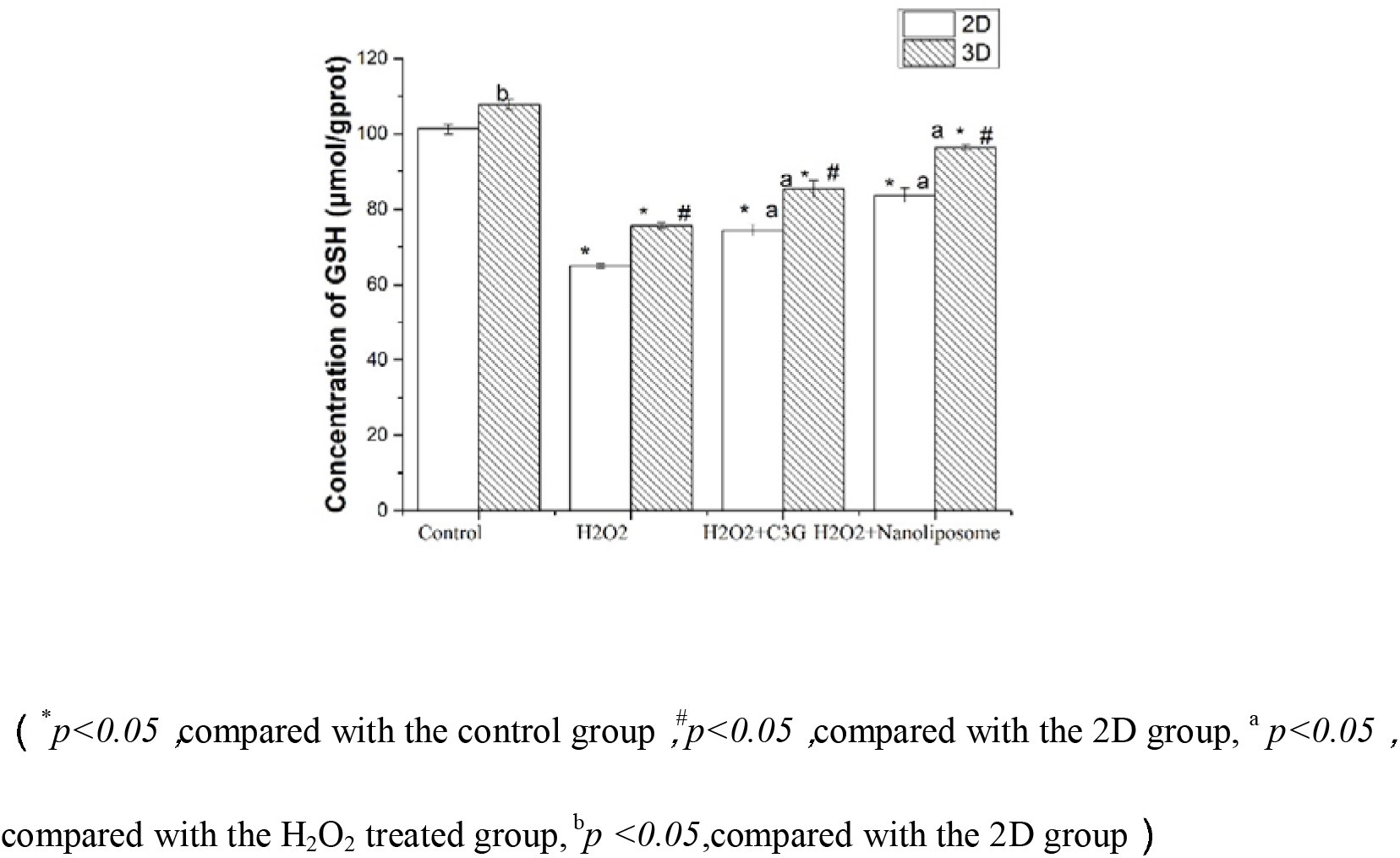
The concentration of GSH in different cell culture models

### 3.5 Assay of SOD activity in different cell culture models

To compare the difference of SOD activity in the 2D and 3D culture models, the Caco-2 cells cultured in the 2D and 3D culture models were treated with H_2_O_2_(0.2mmol), C3G, and C3G liposomes for 12 h, and the SOD activity was tested. Fig. 6 shows the SOD activity in the 2D and 3D cell models, and the SOD activity in different cell culture models is significantly different (*p*<0.05). The C3G and C3G liposomes both can enhance the SOD activity treated with H_2_O_2_, and the enhancement in the 3D model is higher than that in the 2D model. At the same time, the enhancement of C3G liposome is higher than that in C3G.

**Fig. 6.**
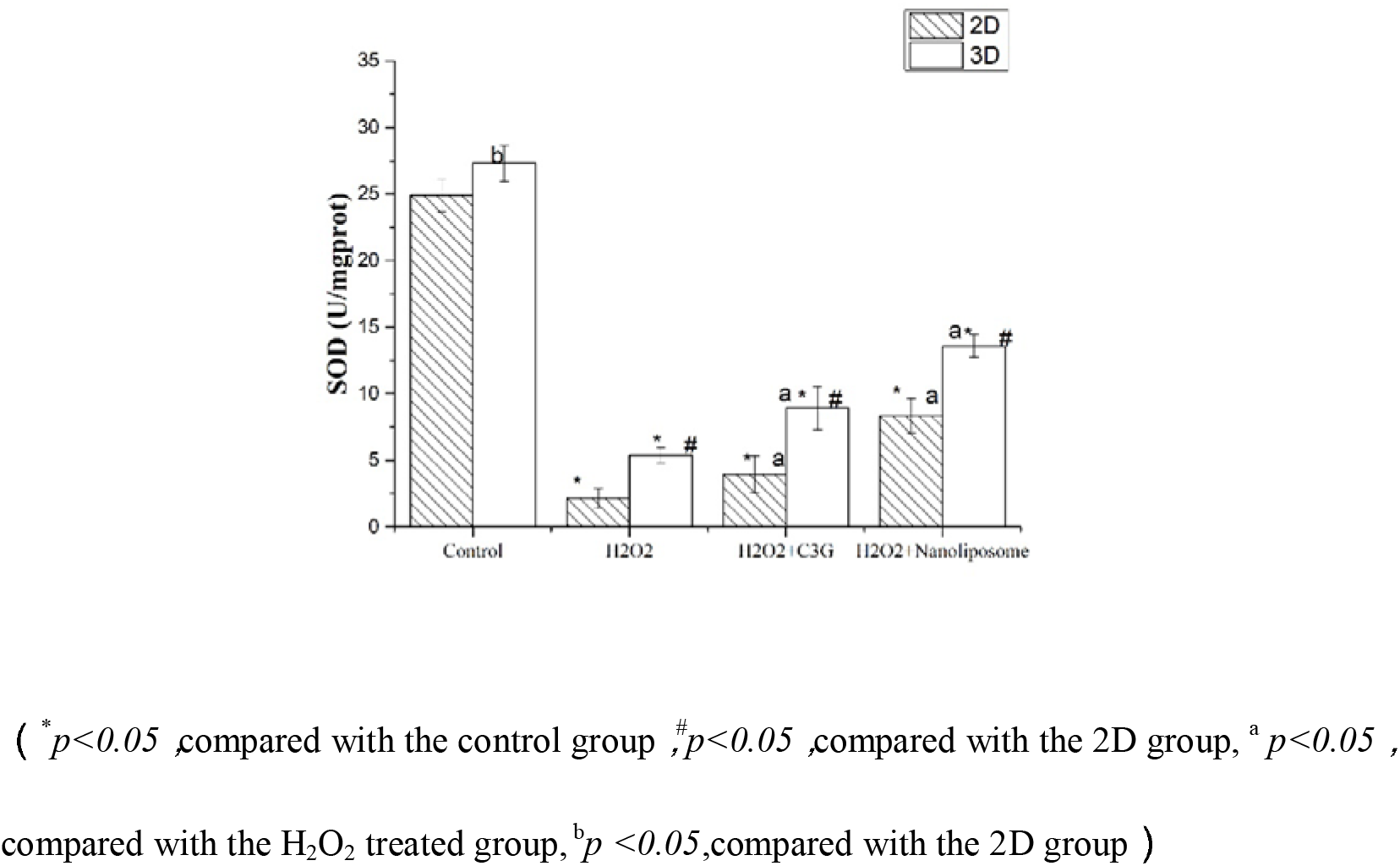
The activity of SOD in different cell culture models

### 3.6 Assay of MDA in different cell culture models

To compare the difference of MDA content in different cell culture models, the Caco-2 cells in different culture models were treated with H_2_O_2_(0.2mmol), C3G, and C3G liposomes for 12 h, and the GSH activity was tested. Figure 7 shows the MDA content in the 2D and 3D cell models, and the result indicated that H_2_O_2_ can increase the MDA content after treatment. The C3G and C3G liposomes both can decrease the MDA content in different cell culture models. As for the reduced effect, the C3G liposomes are higher than C3G. The result showed that C3G and C3G liposomes both can decrease the MDA content in different cell culture models and the reduced effect. The C3G liposomes are higher than C3G. This phenomenon is because the liposomes protect the C3G from the effect of external environment factors, leading to the high effectiveness of cells in different culture models.

**Fig. 7.**
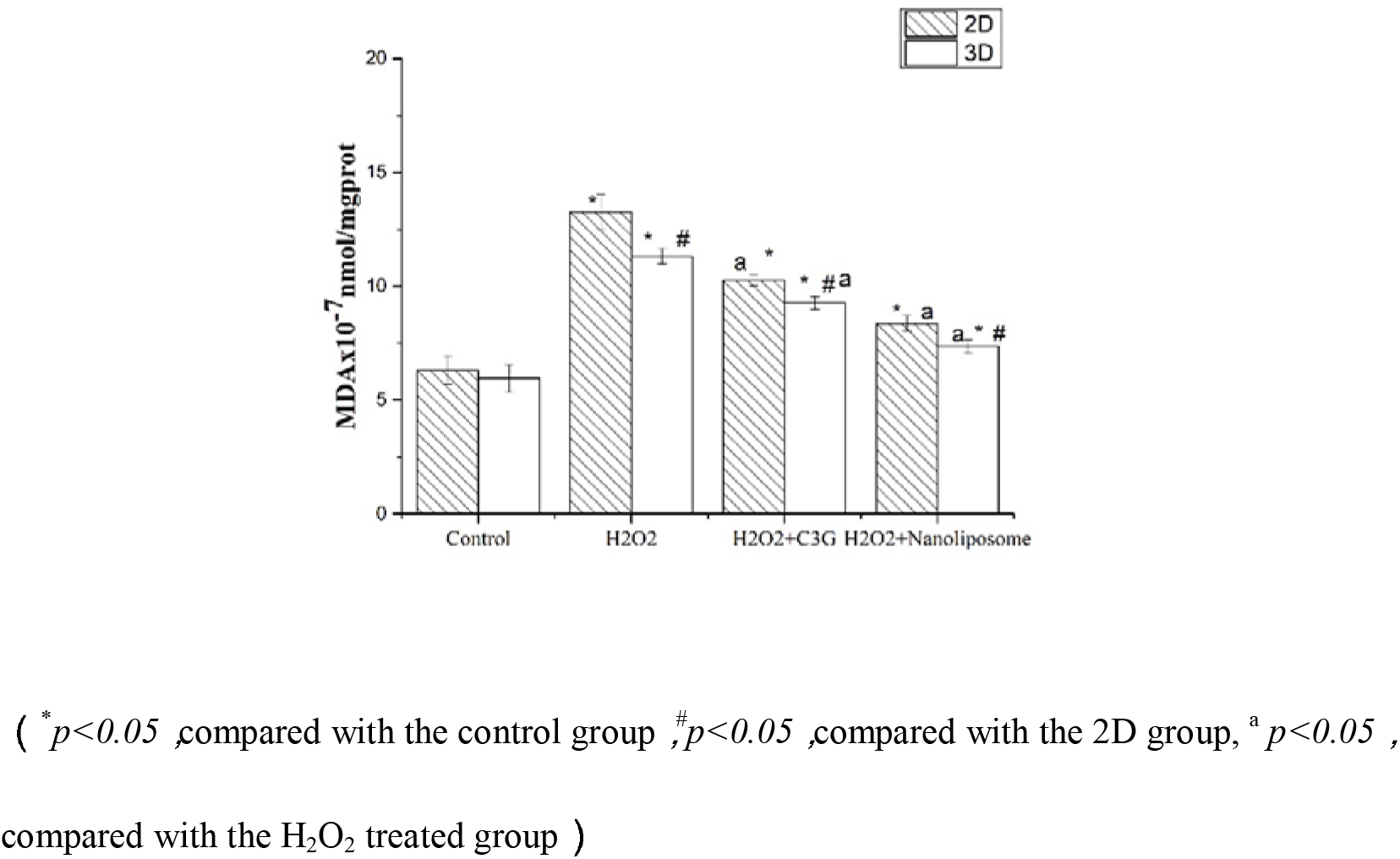
The content of MDA in different cell culture models

### 3.7 Assay of T-AOC in different cell culture models

To compare the difference of T-AOC in different cell culture models, the Caco-2 cells cultured in the 2D and 3D culture models were treated with H_2_O_2_(0.2mmol), C3G, and C3G liposomes for 12 h, and T-AOC assay was tested. Figure 8 shows the T-AOC assay in the 2D and 3D cell models. From Fig. 8, the T-AOC assay in different cell culture models is significantly different (*p*<0.05). The C3G and C3G liposomes both can enhance the assay of T-AOC treated with H_2_O_2_, and the enhancement in the 3D model is higher than that in the 2D model. At the same time, the enhancement of C3G liposome is higher than that of C3G.

**Fig. 8.**
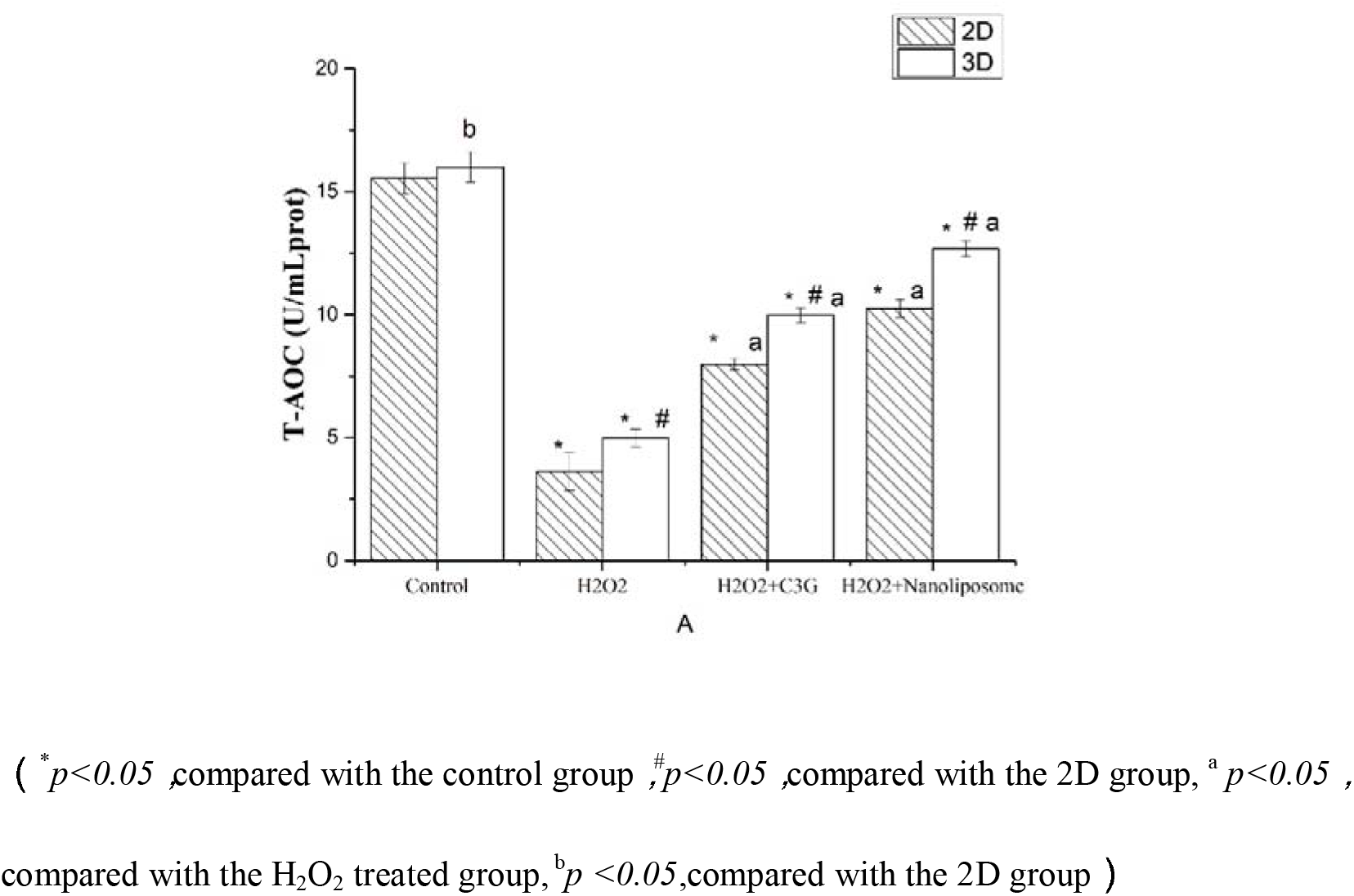
The strength of antioxidant capacity in different cell culture models

The result indicated that the C3G and C3G liposomes can enhance the T-AOC assay and then play an antioxidant role in Caco-2 cells.

## 4. DISCUSSION

In this study, we compared the antioxidant activity of C3G and C3G liposomes in different cell culture models.The cell were treat with H_2_O_2_ and incubated with the C3G and C3G liposomes. The result indicated that : the C3G and C3G liposomes can increase the concentration of GSH, improve the activity of SOD and T-AOC, the content of MDA were decreased after treated. In the 3D cell culture model, the effect is more obvious than cells cultured in 2D model.

C3G is a potent antioxidant that displays anticancer properties in vitro and in vivo^51^, ^52^. However, few studies have evaluated the antioxidant properties of C3G and C3G liposomes in the 3D cell culture model. The results of this study revealed that C3G and C3G liposomes can enhance the activity of GSH, SOD, and T-AOC and decrease the MDA content in Caco-2 cells in certain concentrations. However, this work did not focus on the molecular level of C3G and C3G liposomes to study the antioxidant activity of Caco-2 cells. Several studies have investigated the antioxidant activity of C3G on cells at the molecular level. Thus, further studies on the antioxidant activity of C3G liposomes on cells at the molecular level are needed.

At the same time, the 3D cell culture model was developed as a new in vitro model. The cells cultured in 3D model exhibit features that are close to the complex in vivo conditions^31^ and have been proven realistic for translating the study findings for in vivo applications. In our work, we used hydrogel and built the 3D cell culture model. This method is inconvenient and time consuming. Thus, in future studies, we will use other materials (type I collagen matrix, 3D scaffolds) to construct the 3D cell culture model and compare the differences in different 3D cell culture models. Meanwhile, as for the C3G liposome, a small size of C3G liposomes is expected, making them more effective than C3G. At the same time, the uniform particle size is needed,which can improve the stability of liposomes, because the liposomes were used as carriers, the size is smaller, and they are more effective. The new method, which is uninjurious for liposome preparation, is important in our further studies.

Therefore, we need a new 3D cell culture model to explore the antioxidant mechanisms of C3G and C3G liposomes at the molecular level in the 3D cell culture model. Meanwhile, the properties, expression of specific genes, and protein effects, and mechanisms of C3G liposomes on cells will be explored in further studies, another cells will used to build the 3D cell culture model and study the properties of C3G liposomes in future study is needed.

## 5. CONCLUSIONS

In summary, the mechanism of C3G and C3G liposomes on cells in different cell culture models are as follows. The C3G liposomes and C3G can inhibit cell proliferation. The inhibiting effect of C3G liposomes is higher than that of C3G. In the 3D cell culture model, the inhibiting effect is lower than that in the 2D cell culture model because in 3D conditions, the multicellular resistance and the adhesion between cells reduce the sensibility of Caco-2 cells to the C3G and C3G liposomes. Assay of GSH, SOD, MDA, and T-AOC demonstrated that the C3G and C3G liposomes can enhance the activities of GSH, SOD, and T-AOC and decrease the MDA content in different cell culture models. At the same time, in the 3D cell culture model, these results are more obvious than that in the 2D cell culture model.

## Competing interests

The Author(s) declare(s) that they have no conflicts of interest to disclose.

## Acknowledgements

This work was supported by Zhejiang Provincial Key Laboratory of Biometrology and Inspection and Quarantine, and National & Local United Engineering Lab of Quality Controlling Technology and Instrumentation for Marine Food. We gratefully acknowledge financial support from National Natural Science Foundation of China (31571845), National Key R&D Program of China(grant No.2016YFD0401503), The Key Research and Development Program of Zhejiang province (2018C02049).

## Abbreviations

C3G: Cyanidin-3-O-glucoside
Caco-2: cells Human epithelial colorectal adenocarcinoma cells
MEM: Minimum essential medium
FBS: Fetal Bovine Serum
PC: Phosphatidylcholine
CH: Cholesterol
TEM: Transmission electron microscope
SEM: Scanning electron microscope
2D: Two-dimensional
3D: Three-dimensional

